# Leveraging type 1 diabetes human genetic and genomic data in the T1D Knowledge Portal

**DOI:** 10.1101/2023.02.03.526066

**Authors:** Parul Kudtarkar, Maria C. Costanzo, Ying Sun, Dongkeun Jang, Ryan Koesterer, Josyf C Mychaleckyj, Uma Nayak, Suna Onengut-Gumuscu, Stephen S Rich, Jason A Flannick, Kyle J Gaulton, Noël P Burtt

## Abstract

Translating genetic discoveries for type 1 diabetes (T1D) into mechanistic insight can reveal novel biology and therapeutic targets but remains a major challenge. We developed the T1D Knowledge Portal (T1DKP), a disease-specific resource of genetic and functional annotation data that enables users to develop hypotheses for T1D-based research and target discovery. The T1DKP can be used to query genes and genomic regions for genetic associations, identify epigenomic features, access results of bioinformatic analyses, and obtain expert-curated resources. The T1DKP is available at http://t1d.hugeamp.org.

## Introduction

The etiology of type 1 diabetes (T1D), a complex disease characterized by autoimmune destruction of pancreatic beta cells, is incompletely known (Atkinson, 2012). There are currently no cures or effective prevention strategies, and only recently has an immune intervention (teplizumab) to delay T1D onset been FDA approved. In absence of full blockage of T1D initiation and progression to clinical disease, the only treatment is life-long insulin therapy. There is therefore a pressing need to better understand disease processes as well as to identify new targets for therapeutic intervention. Discoveries from genetic association studies of complex diseases such as T1D can offer novel insight into disease pathogenesis and reveal potential therapeutic targets (Claussnitzer *et al*., 2020), as well as providing human genetic support for pre-existing targets (Dornbos *et al*., 2022).

There are major barriers, however, to translating genetic discoveries into biological and therapeutic insights. The results of genetic association studies are inaccessible to many scientists, since utilizing and interpreting large genetic ‘summary’ files requires expertise in data manipulation and knowledge of domain-specific bioinformatics tools. In addition, most T1D risk variants map to non-coding sequence, where detailed functional annotation of the genome is necessary to predict the affected cell types and genes (ENCODE Project Consortium *et al*., 2020). Finally, testing variant and gene function in cellular and animal models remains a substantial undertaking, often requiring years of work.

Here we report the Type 1 Diabetes Knowledge Portal (T1DKP; https://t1d.hugeamp.org/), an open-access resource developed to help democratize access to genetic, genomic and epigenomic data, and advance T1D research. The primary goal of the T1DKP is to facilitate the generation of accurate, testable hypotheses from genetic association data by providing a user-friendly interface to researchers where they can (i) view the results of analyses integrating genetic and functional annotation data using contemporary bioinformatic tools, (ii) access ‘curated’ resources such as candidate gene lists generated by domain experts in T1D, and (iii) query and visualize data for specific variants, genes, regions, and phenotypes.

## Results

The T1DKP (RRID:SCR_020936) provides human genetic and genomic data relevant to T1D, summarized in **Figure 1**. It currently includes 11 genetic association studies for T1D, including genome-wide association studies (GWAS) from large meta-analyses (Chiou *et al*., 2021), GWAS from biobanks such as FinnGen (https://www.finngen.fi), and targeted, fine-mapping studies using the ImmunoChip (Onengut-Gumuscu *et al*., 2015; Robertson *et al*., 2021). In addition, the T1DKP provides 189 association datasets representing 161 traits relevant to T1D pathophysiology, such as diabetic complications, other autoimmune diseases, and glycemic, lipid, renal, and anthropometric traits.

**Figure 1.**
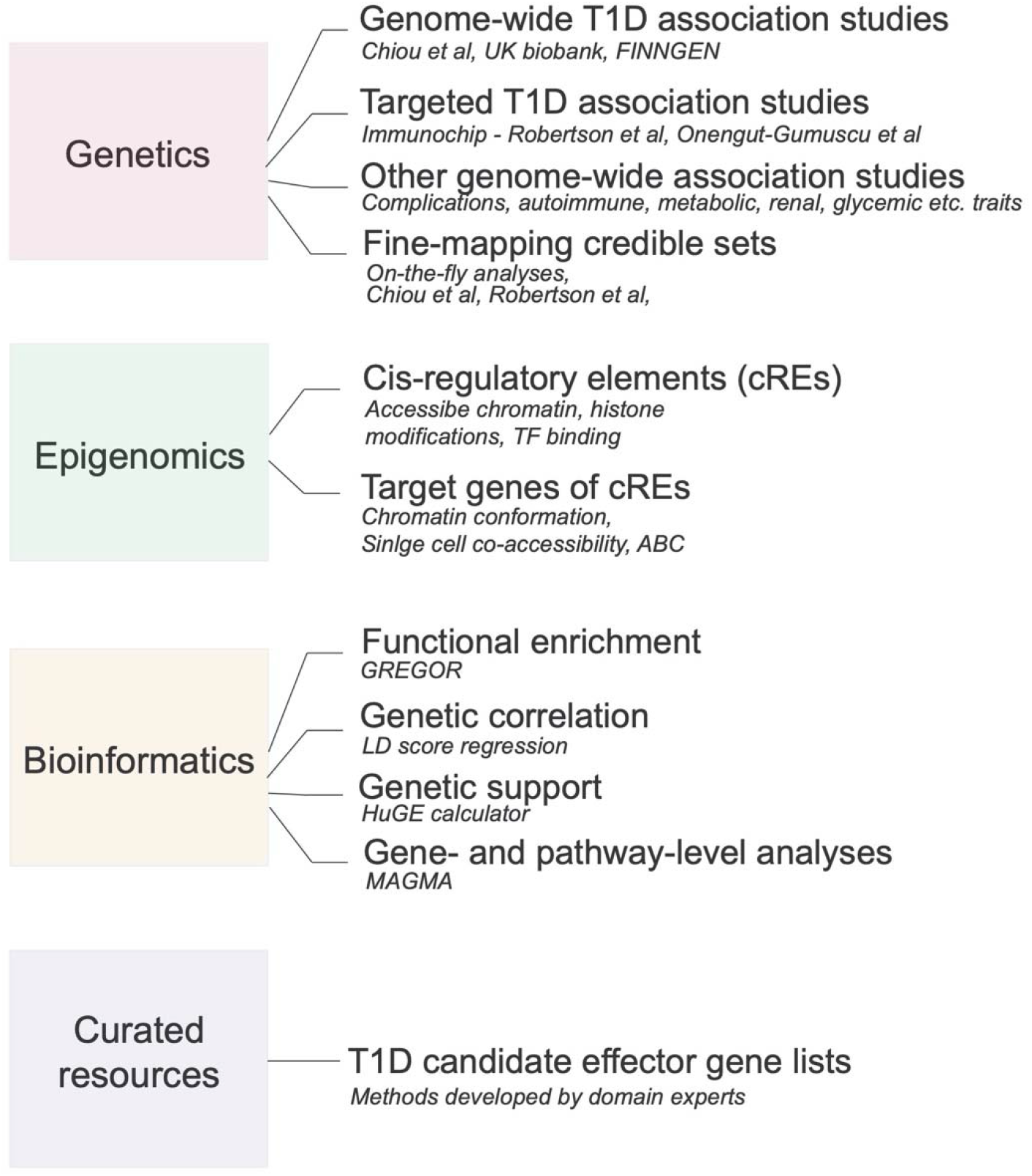
Data content of the T1DKP. The T1DKP provides genetic and genomic data, pre-computed bioinformatics results, and curated resources such as candidate effector gene lists to the T1D community.

The T1DKP aggregates 5,580 functional annotation datasets from the Common Metabolic Diseases Genome Atlas (https://cmdga.org) that describe the location of candidate *cis*-regulatory elements (cCREs) in the human genome and predicted target genes of cCREs in 200 tissues, primary cells, cell lines and stem cell-derived models. These annotations are collected both from resources such as ENCODE (http://www.encodeproject.org) and from genomic studies performed by individual investigators. In the latter case, studies with high value datasets for annotating T1D variants are prioritized for inclusion; for example, there are data identifying cCREs in immune cells in baseline and stimulated conditions (Calderon *et al*., 2019) as well as chromatin interactions linking cCREs to putative target genes in immune cells (Javierre *et al*., 2016).

The T1DKP can be queried to view pages that summarize genetic associations and functional annotations for specific variants, genomic regions, genes, and phenotypes. Visualizations on these pages, such as PheWAS, forest plots, and visualization using the LocusZoom browser (Boughton *et al*., 2021), facilitate user interaction with genetic data. Results from bioinformatic methods integrating genetic and genomic datasets provide additional insight. For example, the gene page includes genetic support analyses that indicate whether the gene is likely involved in a trait (Dornbos *et al*., 2022; de Leeuw *et al*., 2015). In another example, the phenotype page includes analyses that describe functional annotations in different cell types and tissues enriched for trait-associated variants (Schmidt *et al*., 2015), and biological pathways associated with the trait (de Leeuw *et al*., 2015). Several interactive modules can also be accessed from summary pages to enable more detailed investigation. Finally, the T1DKP facilitates independent investigations by providing all genetic and functional annotation datasets for download or programmatic access via a REST API available at http://bioindex.hugeamp.org. Each page and tool of the T1DKP is documented with available online tutorials and videos.

For researchers who are not experts in human genetics (*e*.*g*., trainees, basic biologists, or researchers in an industry setting), the T1DKP offers intuitive distillations of genetic results. On the gene page, the level of genetic support for that gene in T1D is provided by a calculator that assigns a weight from “Compelling” to “No evidence” based on genetic association data (**Figure 2A**). In addition, candidate T1D effector gene lists curated by T1D geneticists are provided, accompanied by evidence supporting each gene, such as protein-coding mutations causing T1D-relevant monogenic phenotypes, non-coding T1D variants linked to the gene, and model system perturbations causing T1D-relevant phenotypes (**Figure 2B**). These lists and supporting evidence can be used to develop hypotheses and guide experiments for specific genes.

**Figure 2.**
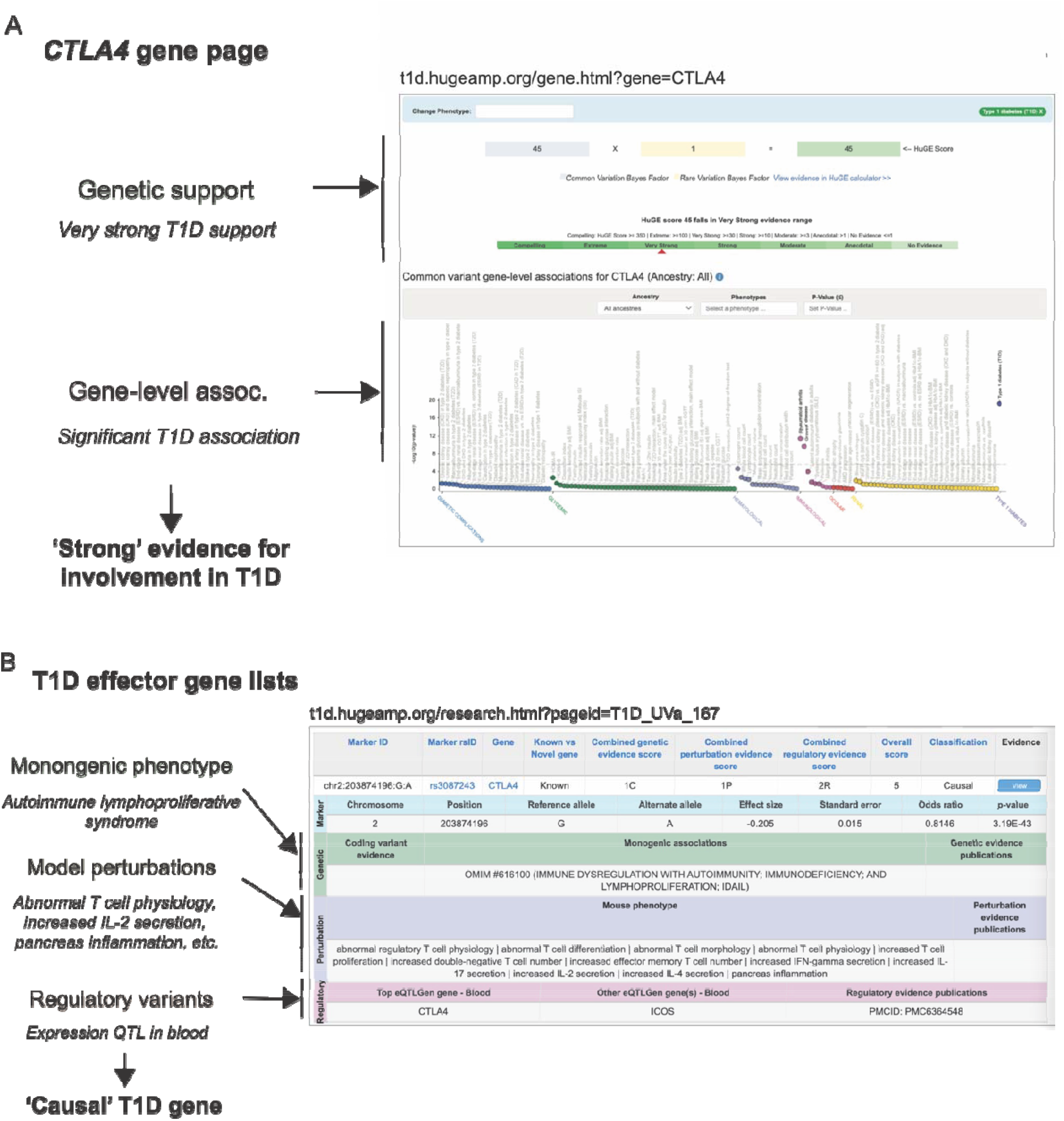
Distilled evidence supporting T1D candidate genes in the T1DKP. The T1DKP provides distillations of human genetic results for researchers who are not experts in human genetics. (A) The summary page for the *CTLA4* gene provides evidence that this gene affects T1D risk, including results providing ‘very strong’ support from the HuGE calculator (Dornbos et al) and strong evidence for T1D association from MAGMA analyses (de Leeuw et al). (B) A ‘T1D effector genes’ list (Onengut-Gumuscu and Rich) predicts *CTLA4* as a ‘causal’ gene for T1D based on genetic, perturbational, and gene regulatory evidence.

For researchers wishing to explore the details of genetic and genomic data in greater detail, the T1DKP provides interfaces and tools that can help to prioritize candidate genes likely involved in T1D risk at specific loci. For example, from the region page the user can link to a ‘Variant Sifter’ module which enables selection of a series of filters to prioritize candidate variants, genes, and tissues/cell types to guide experiments in that region (**Figure 3**).

**Figure 3.**
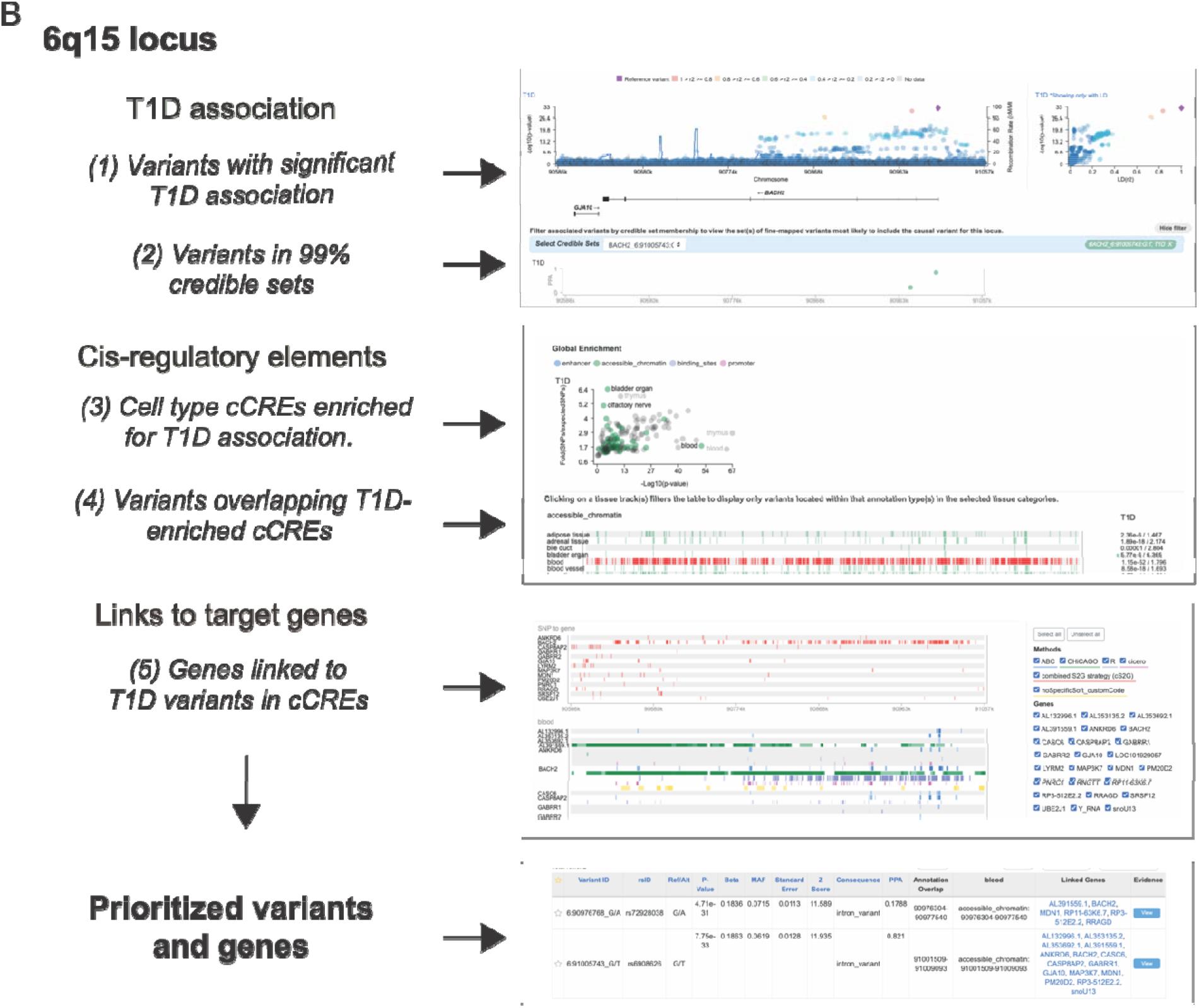
Predicting causal mechanisms at T1D risk loci in the T1DKP. Predicting causal mechanisms at the 6q15 locus. (top) Prioritizing variants with evidence for affecting T1D risk based on significant association and 99% credible sets. (middle) Prioritizing variants overlapping cCREs active in T1D-enriched cell types and tissues. (bottom) Prioritizing genes linked to variants in cCREs in specific cell types and tissues. From these analyses two variants are predicted as causal candidates for T1D risk at this locus, which are linked to multiple candidate genes including *BACH2* in immune cells.

## Discussion

The T1DKP enables exploration of genetic and functional annotation data relevant to T1D on an interactive website that is useful to both the experimental biologist and the expert in human genetics. Compared to disease-agnostic resources that also provide platforms for analyzing human genetic and genomic data such as Open Targets (opentargets.org), two core strengths of a disease-focused resource such as T1DKP are 1) the aggregation of data sets from studies of high value to that specific disease that may be missing from ‘pan-disease’ catalogs and 2) the incorporation of curated datasets created by domain experts. Moving forward, a key goal of the T1DKP is to continue engaging with the T1D community to identify and add T1D-specific datasets.

In the future, the T1DKP will incorporate different types of genetic and genomic data that provide additional value for hypothesis generation beyond currently available data. For example, genetic association data from whole genome and exome sequencing studies of T1D cohorts will help identify genes carrying rare risk variants involved in T1D. Similarly, genomic data from populations of diverse genetic ancestry as well as systematic screens of variant function will allow more comprehensive prioritization of risk loci. Finally, incorporating data on gene perturbation phenotypes in human cells and model organisms will facilitate prioritizing genes involved in T1D. We look forward to collaborating with the T1D community to advance these and other areas of the T1DKP.

## Acknowledgements

This work was supported by the National Institutes of Health grant numbers DK105554 to JF, NB and KG, DK122607 to KG, DK122586 to SR. We thank members of the Gaulton, Burtt, Flannick, and Rich labs for input on the manuscript, and members of the AMP-T2D and AMP-CMD consortia and the T1D research community for critical input on the development of the T1DKP and other repositories in the Knowledge Portal Network.

